# Effect of habitat quality, microclimatic conditions and waste water contamination on diversity & distribution of Collembola community

**DOI:** 10.1101/668749

**Authors:** Mohammad Jalaluddin Abbas, Hina Parwez

## Abstract

Soil biodiversity is undergoing dramatic changes due to human and climatic interference, habitat loss and fragmentation due to land use change, increasing pollution and degradation in terrestrial ecosystems. The overall effect of habitat loss is on the loss of soil biodiversity which might be due to negative response of substrate quality. Adverse microclimatic conditions may also be detrimental to species specific communities along with waste water contamination. Thus, an investigation has been carried out to ascertain the effect of habitat quality, microclimatic conditions and waste water contamination on diversity and distribution of Collembola community at Aligarh. For this, 4 samples have been taken monthly for a period of one year from two different study sites of Aligarh along with edapho-chemical properties. We observed that, density of Collembola (18.8 ind./sample) was always approximately three times more in agriculture site than in wasteland (7.6 ind./sample) site. Population of Collembola was statistically positive significant with reference to soil organic matter % (r = 0.621, P>0.05), SOC % (r = 0.622, P>0.05), Available nitrogen (r = 0.622, P>0.05) and Potash (r = 0.562, P>0.05) in Agriculture site whereas these all were statistically insignificant in Wasteland site. This observation suggests that quality of habitat has a direct effect on the diversity and population of Collembolans. The wasteland site which is a sewage filled from the nearby colonies also it is a dumping ground of waste building material under Municipal Corporation. Hence, the soil is highly contaminated as compared to the agricultural soil.

## Introduction

Human activities tends to decrease the abundance and diversity of euedaphic springtails including soil compaction through trembling (Massoud et al. 1984), Vehicular activity (Aritajat et al. 1977, Heisler 1991 and 1994, Heisler and Kaiser 1995) and urbanization (Pouyat et al. 1994, Sterzynska 1989). On the other hand, habitat feature that might seem to have the most influence on Collembola is vegetation (Hopkin 1997). Thus, Beng and Bengtsson (2007), suggested that substrate quality mostly determines the distribution of soil microarthropods in the soil profile. Sjursen, Michelsen and Jonasson (2005) have stated earlier that soil microarthropods are bottom up controlled probably linked to changes in food quality and their availability rather than by direct climatic influences. Although, substrate quality may be the main driver which affects soil microarthropods distribution as well as their densities and ultimately abundance may varied in varied manner. However, several numerous ecologists have concluded that urbanization may have several effects on soil invertebrate communities (Santorufo et al. 2012) thus, species diversity and abundance decreases (Battigelli and Marshall 1993, Eitminaviciute 2006, and Gongalsky et al. 2010).

Interestingly, soil quality is an important factor affects the soil microarthropods communities in terms of their diversity loss as well as on their fecundity. Apart from soil inhabiting microarthropods communities, Collembolans are good bio-indicators of arable soil quality (Rusek-1998), are very sensitive to variations in soil environment (Parisi-2001) and conditioning to detritus for microbial breakdown as well as farming soil to form soil micro-structure (Chahartaghi-2005). However, Collembola communities are still poorly understood (Alvarez et al. 2000) in spite of the role of these soil organisms in mineralization and humification of soil organic matter (Coleman 1985, Huhta et al. 1988, Czarnecki 1989, Striganova 1992) and they are considered as bio-indicators in studies of soil quality (Heisler 1995, Kopeszki 1997). This is not yet surprising fact that, activities of all soil born organisms and processes are affected by levels of soil organic carbon, moisture contents in soil and temperature regimes in a terrestrial environment which are dependent on variety of other soil physio-chemical factors, environmental factors and vegetation cover.

There is a paucity of information from agricultural soils of North India region in relation to the density and diversity patterns of Collembolans except very few studies reported soil microarthropods diversity from West Indian soils such as West Bengal (Chaudhury and Roy-1967, 1971a, 1972, Hazra and Chaudhury-1990, Chaudhury et al. 1978, Hazra-1978a and 1978b and Mitra et al. 1977, 1981, 1983). However, no study has been reported up to till date as a comparative study between an arable land and urban or degraded soil from Indian continent including this area. Thus, aims of this study have been both quality and quantity based comparisons with in two different study sites which are more sensitive to diversity of soil microarthropods as well as in terms of different management systems. Among these study sites, one has to expect a precursor to diversity of soil microarthropods (arable land management site) and other (wasteland management site) has to detrimental for diversity of soil microarthropods.

## Materials and Methods

### Area of study

The area selected for study is situated at Aligarh. It is a flat topographical area, located in western part of UP at latitude 27-54’N, longitude 78-05E’ and altitude 187.45 meter above sea level. It is a subtropical zone with fluctuating climatic conditions consisting of four different seasons characterized by extreme winter and summer followed by medium to heavy rainfall during monsoon months and a post monsoon, sweet spring. In hot dry summer, the temperature rises up to 48 ºC, while in winter cold, the temperature goes drops up to 2 ºC. Relative humidity also fluctuates with the sudden change of environmental temperature and with rainfall patterns. Such widely varying climatic conditions provide a variety of ecological niche to soil dwelling animals and interesting for ecological studies on soil microarthropods in this region.

#### (a) Study Sites

Two sites have been selected for dynamics and ecological study of Collembola community-

Site (I) Agricultural land

Site (II) Wasteland site

##### Site (I) Agricultural land

An agricultural site was selected at Pangipur village that is very close to University Qila which is situated at outskirts of Aligarh city. The area of selected agricultural field was approximately 2 acre. Various crops were harvested with in a year such as, Mung been (*Vigna radiata* L.), Arhar (*Cajanus cajan*) and Wheat (*Triticum aestivum*) etc. The site was well managed by its farmer in terms of tilling, irrigation and fertilizers used time to time in the field. The soil of site was alluvial type, a mixture of sand, silt and clay.

##### Site (II) Wasteland site

This area was selected at wasteland site as it is confirm with its famous name known as Lal Diggi, and samples have been collected from around the Lal Diggi pond. The pond site was polluted by flowing sewage water comes from nearby residential houses of its civilian area. A cross road is also linked on the two side of this pond so that, vehicular disturbance also the source of air pollution as recorded in the present study.

#### (b) Sampling and extraction of Collembola

As mentioned earlier, samples were taken every week regularly and the points selected within the plots were distributed randomly. Total, 96 samples have been taken for site study during the investigation period from both sites. Each sample consists of 4 corers of 5 cm. size. Modified Tullgren funnel apparatus was used for extraction of Collembola. Collembolans were collected inside a beaker which contained 70 % alcohol with few drops of glycerol. Before permanently mounting of soil microarthropods, they were preserved in ethanol alcohol via series of alcohol (70%→ 80%→ 90%) having each of one series for an hour of time period and finally 10-20 minutes preserved in 100% alcohol. This method was actually made because of clarity of soil faunal material. After preservation, they were separated and mounted with DPX.

#### (c) Identification

Collembolans were identified up to the level of their family, genera or species as possible using a range of taxonomic keys. A binocular stereomicroscope (OLYMPUS, CX-21) was used to identify them.

#### (d) Analysis of edaphic and chemical parameters

Temperature of the soil was measured by directly inserting the thermometer into the soil up to the required soil depth and Relative Humidity was determined by Dial hydrometer. Soil Moisture has been determined by using the method as described by Dowdswell (1959). Soil pH, Organic Carbon, and Nitrogen, all were examined by standard laboratory methods.

#### (e) Statistical Analysis

To study the population dynamics of Collembolans, the parameters considered were density, abundance, fractional population, relative density, and absolute frequency between two different quality habitats. Pearson’s regression tests were performed to evaluate the relationships between Collembola population and each soil physical and chemical characteristic. One way analysis of variance (ANOVA) was calculated with the help of SPSS package (SPSS 18.0 Version) to evaluate the correlation between populations of Collembola community with reference to each physical and chemical parameter.

## Results

### (a) Results pertaining to population dynamics of Collembola

Total 4,729 individuals have been collected during study in which 2,731 individuals (57.7 %) from agriculture site and remaining 1,998 individuals (43.3 %) from wasteland site. Thus, compared to wasteland site, more than 14 % microarthropods population was recorded in agriculture site. More than half of the specimens were Collembolans recorded in agriculture site. The proportion of Collembola in samples from each habitat type varied considerably as listed 57.8% in agriculture site and 25.6% in wasteland site. Densities of Collembola (18.75 ind/samples) was always approximately three times more in agriculture site than in wasteland (7.60 ind/samples) site, where in case of abundance, Collembolans were frequently abundant in agriculture site (19.96 ind/samples) than wasteland site (8.07 ind/samples) as shown in table 1. The peak population of Collembolans buildup in post monsoon (winter and spring altogether) months in both study sites as shown in figure-1.

**Table 1.** Density and abundance of Collembola population in two different habitats of Aligarh

| Time |  | Agriculture site |  | Wasteland Site |  |
| --- | --- | --- | --- | --- | --- |
|  |  | Density | Abundance | Density | Abundance |
| Apr-10 | Summer | 5.00 | 6.67 | 4.00 | 4.00 |
| May |  | 4.00 | 5.33 | 1.25 | 2.50 |
| June |  | --- | --- | 0.75 | 3.00 |
| July | Fall | 2.50 | 5.00 | 8.50 | 8.50 |
| Aug. |  | 10.00 | 10.00 | 9.00 | 9.00 |
| Sept. |  | 7.00 | 9.33 | 2.75 | 3.67 |
| Oct. | Winter | 4.00 | 8.00 | 3.75 | 5.00 |
| Nov. |  | 6.50 | 8.67 | 7.50 | 7.50 |
| Dec. |  | 34.50 | 34.50 | 6.50 | 6.50 |
| Jan. 11 | Spring | 62.25 | 62.25 | 19.25 | 19.25 |
| Feb. |  | 70.25 | 70.25 | 20.50 | 20.50 |
| Mar. |  | 19.50 | 19.50 | 7.50 | 7.50 |
| Average |  | 18.75 | 19.96 | 7.60 | 8.07 |

**Figure 1.**
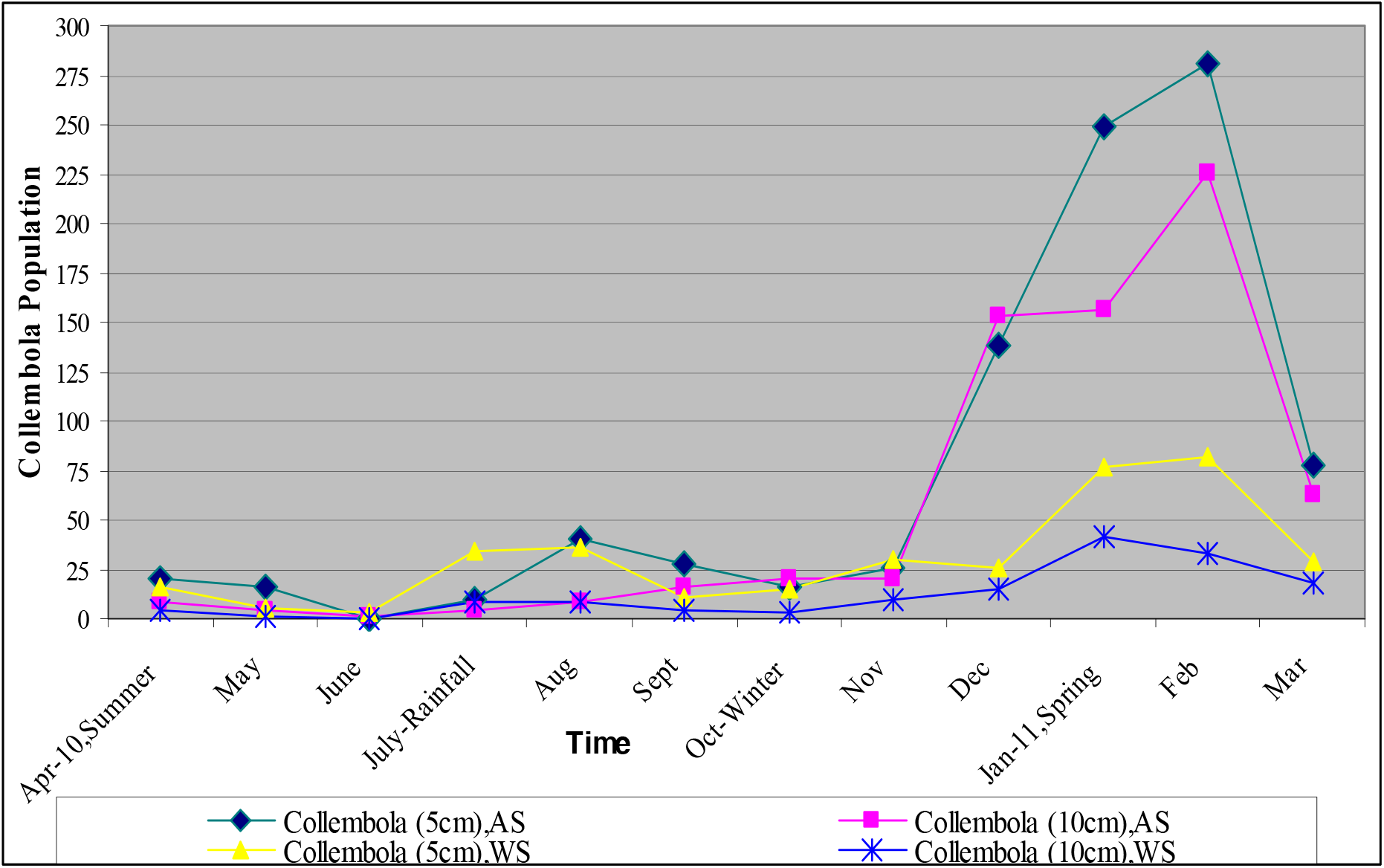
Temporal dynamics of Collembola community in two different sites and depths

### (b) Results pertaining to Correlation between soil matrix and Collembola population

Both soil temperature (r = −0.794, P<0.05) and pH (r = −0.848, P<0.05) both were statistically negative significant with reference to Collembola population in agriculture site. The population of Collembola was statistically negative significant (r = −0.749, P<0.05) with reference to soil temperature whereas positively significant with reference to potash (r = 0.675, P>0.05) and phosphate (r = 0.619, P>0.05) in wasteland site (table-3A). Population of Collembola was statistically positively significant with reference to SOM % (r = 0.621, P>0.05), SOC % (r = 0.622, P>0.05), Available nitrogen (r = 0.622, P>0.05) and potash (r = 0.562, P>0.05) in agriculture site (table 3A).

The effects of microclimatic conditions on population, density and abundance of soil microarthropods may be due the varied nature of functional interactions in a particularly different terrestrial ecosystem that may dependent on soil quality as it correspondence soil matrix such as concentration of soil organic carbon, total nitrogen available (figure 2A and 2B) and C/N ratio showed the distinct temporal changes in pedoecosystem of tropical climatic region of Uttar Pradesh.

The more significant difference in chemical and physical parameters has shown in table-2 with reference to Collembola population in both study sites. Among the soil chemical parameters, soil nitrogen organic carbon contents and soil organic matter, all have significant positive correlation with reference to Collembola population in arable soil whereas statistically insignificant in wasteland site. This is remarkable difference between both study sites. Hence, it can be said that, arable soil condition and its management are better to growth and support of population buildup of Collembola whereas totally different conditions in wasteland site that may counterintuitive for faunal population survival in this site. In some extent it was duly viewed that potash and phosphate contents, both were positively significant correlated with respect to Collembola population in wasteland site. This may precursor to soil faunal life in waste water contaminated site however the reason of its nature is not clearly understood.

**Table 2.** Correlation between physio-chemical properties of soil and Collembola population in two different habitats of Aligarh

| Parameters | Agriculture site | Wasteland Site |
| --- | --- | --- |
|  | r-value | r-value |
| Soil temperature(°C) | -0.794* | -0.749* |
| Soil moisture (%) | 0.193 | 0.195 |
| Relative Humidity (%) | 0.622* | 0.377 |
| Soil pH | -0.848* | 0.074 |
| SOM (%) | 0.621* | -0.013 |
| SOC (%) | 0.622* | 0.002 |
| Available N (ppm) | 0.622* | 0.002 |
| Potash (ppm) | 0.562* | 0.765* |
| Phosphate (kg/acr) | -0.046 | 0.619* |

(\*values are significant at $p > 0.05$ )

### (c) Seasonal abundance of Collembola population

The test of within subject effects and between subject effects for season and its interaction with population growth indicated the significant amount of abundance of Collembola in both study sites (tables-1). Thus, seasonal fluctuation in population of Collembola may exhibited more or less but similar pattern of changes in both study sites. Under the priority of data observed from both study sites, one thing is clear that January and February were frequently abundant months followed by a small peak in August (table-1). This is due to perhaps the sudden decline of temperature regimes in August and more stable spring environment (moderate temperature and moisture conditions) in January and February months.

The soil Collembola population slightly declined in April and May with least abundant in June. This showed a subsequent decline in population of Collembola due to the fact that temperature of the environment as well as soil increased sharply along with Relative humidity through April to June. Therefore, the faunal population decreased gradually up to July. Samiuddin and Haider (1999) also demonstrated a relation between population fluctuation of soil microarthropods and soil temperature. They have stated that, soil temperature as one of the main factor influencing the population fluctuation of soil microarthropods. Other than this study in Indian continent, Tripathi et al. (2007) stated that, edaphic factors directly influence the soil faunal abundance.

The less affected collembolan community in rainy months (fall season) in wasteland site due to the fact that, rainfall is the key component that influence the collembolan population in terrestrial ecosystems. Far less this observation, more than 70 % effect has been observed during spring season that was the more suitable time for high density development of collembolan communities (table 1).

Prior to this investigation, there was a remarkable difference in population composition of Collembola vertically in both study sites. The vertical distribution patterns however were same in both study sites, but more individual population of Collembola was found in 0-5cm depth as compare to 5-10cm deep soil (figure-1).

## Discussion

To conserve biodiversity in increasing fragmented landscapes requires the better management strategies because we should not only understand the response of species and communities but also the effects of changing habitat quality and habitat network of each community structure. Urbanization may have several effects on soil invertebrate communities in general as well as in particular on a single species. Furthermore, understanding the multiple effects of habitat quality on community dynamics has been limited because it is frequently less investigated. However, general trend is that, quality of soil directly affects the density and abundance of soil fauna as we also observed in this study of Collembola diversity loss in low soil quality habitat.

Although, the investigated soils were collected in the same urban areas however, their chemical and physical properties of soil strongly differed in both study sites. The lower densities of soil inhabitants tend to decrease the quality of soil some where in food resources, soil microstructure and microclimatic conditions derived by harsh climatic conditions, pressures and threats, in flooding waste water polluted areas and due to air pollution in nearby urban systems. This is because of, Urban ecosystems are characterized by high density human habitation and intense transport processes those all are detrimental to soil organisms. Thus, urbanization comes of several forms of disturbances, changes of temperature, pH and moisture regimes, edaphic conditions and pollution (Cristina Fiera 2009).

Climatic factors belong to the most determinant ones. According to the findings of present research study, we have concluded that, there were marked variations between seasons and population of each community during the year. Thus, seasonal patterns are responsible for population dynamics, diversity, abundance and ultimately the structure of each community. However, the advantages of coupling reproduction with the most probable moist period may be related to quantity and quality of food resources. Most of the food resources were critically less distributed in wasteland site that they do not response to influence the diversity of Collembola communities. This may be also due to low habitat quality that directly affects the diversity niche of Collembola communities. Thus, local habitat conditions or its quality may serve as a strong indicator of changing diversity patterns. Our observations clearly indicate about this important phenomenon. In addition to this phenomenon, Sjursen, Michelsen and Jonasson (2005) have proposed that soil microarthropods are bottom up controlled probably linked to changes in food quality and their availability rather than by direct climatic influences.

In arable soils, the soil microarthropod community is rather simple, but it may become complex in contaminated or polluted soils where more suitable conditions may not favorable for colonization and further settlement of most of microarthropod communities. Various components such as soil moisture, temperature and availability of energy source (carbon) determine the activity of the soil organisms. Moisture, temperature and carbon availability need to be synchronized for optimal performance of a range of key biological functions in different habitats. However, these all are time dependent or may be seasonal because of the change of the diurnal temperature which may affect directly on edaphic conditions of soil ultimately change the diversity of Collembolans. Thus, it may hypothesize that, **temperature and soil moisture both are precursor of Collembolans diversity whereas relative humidity is the controversy between these edaphic factors**.

The continued drainage of contaminated water may lead to accumulation of heavy metals in soils and adverse effects on soil living biological communities. Most species of Acari (mites) are higher accumulators whereas Collembolans are poor accumulators. Collembolans were less abundant in wasteland site which was highly polluted by waste water and highly abundant in agriculture site that was irrigated by clean potable water. Thus, the findings of this study clearly indicate that, among soil microarthropods, Collembolans have a high potential as bio-indicators of environmental changes including pollution of soils in India. In view of this observation, it can be hypothesized that, **the numerical response of soil microarthropods may varied considerably in different quality habitats of soil** as we recorded in our study.

Climate change is an ultimate source of changing diversity in terrestrial ecosystems because we have also observed in our study that temperature and soil moisture have cumulative affect along with relative humidity on the diversity patterns of soil microarthropods as stated earlier. Direct effect of Ultra Violet Radiation increases the temperature of a particular terrestrial ecosystem. Thus, temperature regimes are most effective in growth and survival as well as reproduction of Collembola communities.

Collembolans are susceptible to local disturbances mostly in urban soils and rarely in farming systems however, population of Collembola community directly or indirectly affected by the temporal and spatial scaling of pollutant distributed or, resistant to soil matrix and microclimatic conditions. The relative scaling of pollutant distributed in wasteland site was not understood clearly because of highly heterogenous soil environment due to contaminated waste water. (Forbes and Kure 1997) also stated in this line that, local extinctions may be created by some toxic effects of any pollutant or contamination agent and the effect dynamic comes out if the sediment and water processing activities of the resident organisms are influenced by expose levels that they have created (Forbes and Kure 1997).

There is evidence that soil Collembolans shows a density dependence dispersal rate that may be influenced by food density and moulting intensity (Bengtsson et al. 1994 a). Young adults seem to disperse more readily than both older ones and juveniles. This is due to perhaps reflecting a risk spreading behavior in matured Collembolans, who lay several separate egg batches during their life time (Bengtsson et al. 1985).On the other hand, reproduction, moulting intensity and growth of individuals of Collembola community are mostly depend on physical and chemical characteristics of soil. Thus, habitat quality may results the classical pattern of density for each species specific or community level along with suitability of its environment. Because the breakthrough of species specific population of Collembola community is totally time dependent. However, density rate may vary from one habitat to another because of flexibility of microclimatic conditions in different habitats at one regional level. In this condition, the probability is that, high density may be found in good quality soils, which have suitable food resources and limited climatic interference.

### Future lines of investigation based on the outcome of this study

The source of pollution however clearly understood as it was waste water comes from nearby civilian area which could be consider as indicator of diversity loss however to find out more clear evidence, much more investigation are required from a randomly selected (wasteland) site to tell whether Collembola density will influence on which condition and how they could recolonize for further settlements.

## Conclusions

Urban sites are often harsh and characterized by many pressure and threats from limited growing space to adverse climatic conditions and air pollution (Cristina Fiera 2009) along with water contamination. This is due to explosive human population in urban areas and due to lower class management strategies for well being of both city dwellers and nature. Thus, there is an urgent need to develop a simple protocol to assess the effect of drainage pollution and waste water contamination that is an ultimate source of soil degradation because it is much necessary to conserve native soil microarthroods biodiversity and to minimize the adverse effects.

Collembolans showed a consistent trend of fluctuations with time as a function of climate. However, the factors like soil moisture, temperature, relative humidity, pH of soil, and food resources, all have somewhere cumulative effect in a varied manner along with quality of soil preferably. This is certainly most important requirement to conserve the diversity loss in polluted land sites. Thus, our management should be more appreciable than our frequently using policies. It requires the uniformity in all along the residential areas and for conserve the nature’s invisible currencies. This is because of flooding areas can cause the dramatic loss of soil diversity that is highly effective to survival of soil microarthropods.

Climatic manipulations are also responsible for survival of soil faunal diversity. Thus, seasonal patterns are another important indicator for population buildup of Collembolans. In the light of observations of this study, it may concluded that cool temperatures with medium humid environment in winter and spring months are best possible environmental conditions for survival and rich diversity of Collembolans. Farmers may use these situations and conditions to get more profit in terms of highly productive manner of these seasons. Because of the diverse functional activities in these seasons may influence the growth and support of perennial crop plants as a result of high productivity due to maximum diversity as well as functional activities of soil microarthropods.

## References

Alvarez T., Frampton G.K., and Goulson D. (2000).The role of hedgerows in the decolonization of arable fields by epigeal Collembola. Pedobiologia, 44; 516–526.

Chahartaghi M. (2005). Feeding guilds in Collembola based on nitrogen stable isotope ratios. Soil Biology and Biochemistry. 37; 1718–1725.

Chaudhuri D.K. and Roy S.(1972). An ecological study on Collembola of West Bengal, India. Ric. Indian Museum India. 66 (1-4); 81–101.

Chaudhuri D.K. and Roy S. (1971a). Interaction between soil Collembola and other subterranean arthropods. Sci. Cult. Calcutta. 36(5); 280–282.

Chaudhury D.K. and Roy S. (1967). Qualitative composition of Collembola fauna of some uncultivated fields in Nadia district (West Bengal) with correlation between monthly population and individual soil factors. Rev. Ecol. Biol. Soil. 4(3); 507–515.

Chaudhury D.K., Hazra A.K. and Roys S. (1978). Soil factors governing the distribution of Collembolans (Insecta) in the graveyard of Berhampore, Murshidabad district (West Bengal). In: Soil Biology and Ecology, India (Ed. C.A. Edwards and G.K. Veeresh) UAS Tech. Series No. 22; 161–172.

Coleman D.C. (1985). Through a ped darkly: An ecological assessment of root-soil-microbial-faunal interactions. (In: Ecological Interactions in Soil. Eds: A.H. Fitter, D. Atkinson, D.J. Read and M.B. Usher) –Special Publ. NO.4. Br. Ecol. Soc; 1–21.

Coleman D.C., Crossley D.A. and Hendrix P.E. (2004). Fundamentals of soil ecology, second edition. Academic Press, San Diego.

Czarnecki A. (1989). Collembolans jako element biologicznego systemu na obszarach podlegajacych silnej antropopresji [Collembola as a component of biological system of the areas under strong anthropopresion] Habilitation Thesis – UMK, Torun 156pp. (in Polish).

Dowd Swell W.H. (1959). Practical Animal Ecology. Methuen and Co. Ltd. London.

Hazra A.K. (1978a). Effects of organic matter and water content of soil on the distribution of Collembolans (Insecta) in an uncultivated field of West Bengal. Bull. Zool. Surv. India. 1(2); 107–114.

Hazra A.K. (1978b). Ecology of Collembola in a deciduous forest floor of Birbhum district, West Bengal in relation to soil moisture. Oriental Ins. 12(2); 265–274.

Hazra A.K. and Chaudhury D.K. (1990). Ecology of soil arthropod fauna. Rec. Zool. Surv. India, Occ. Paper No. 120; 1–295.

Heisler C. (1995). Collembolans and Gamasina –bioindicators for soil compaction – Acta Zool. Fennica. 196; 200–205.

Huhta V., Setala H. and Haimi J. (1988). Leaching of N and C from beach leaf litter and raw humus with special emphasis on the influence of soil fauna – Soil Biol. Biochem. 20; 875–878.

Kopeszki H. (1997). An active bioindication method for the diagnosis of soil properties using Collembola – Pedobiologia. 44; 159–166.

Mitra S.K., Dutta A.L., Mondal S.H., and Sengupta D. (1983). Preliminari observation on the effect of rotation of crops and fertilizers on Collembola. New Trends in Soil Biology. Ed. by P. Lebrun. 657–663.

Mitra S.K., Hazra A.K. and Sanyal A.K. (1977). Ecology of Collembola at Eden Gardens, Calcutta. Ecol. Bull. (Stockholm). 25; 539–544.

Mitra S.K., Hazra A.K. and Sanyal A.K. and Mondal S.B. (1981). Changes in the population structure of Collembola and Acarina in a grassland ecosystem at Calcutta. In: G.K. Veeresh (ed.). Progress in Soil Biology and Ecology in India. UAS Tech. Series No. 37; 143–146.

Parisi V. (2001). The biological soil quality, a method based on microarthropods. Acta Naturalica de L’ Ateneo Parmense. 37; 97–106 (in Italian).

Rusek J. (1998). Biodiversity of Collembola and their functional role in the ecosystem. Biodiversity and Conservation. 7; 1207–1219.

Striganova B.R. (1992). Trophic relations in soil animal communities and decomposition rates. Pol. Ecol. Stud. 16; 119–130.

